# Gut microbiota impairs insulin clearance during obesity

**DOI:** 10.1101/2020.05.08.083923

**Authors:** Kevin P. Foley, Soumaya Zlitni, Brittany M. Duggan, Nicole G. Barra, Fernando F. Anhê, Joseph F. Cavallari, Brandyn D. Henriksbo, Cassandra Y. Chen, Michael Huang, Trevor C. Lau, Jonathan D. Schertzer

**Author notes:** Corresponding author: Dr. Jonathan D. Schertzer, Department of Biochemistry and Biomedical Sciences, Faculty of Health Sciences, McMaster University, HSC 4H30D; 1200 Main Street West, Hamilton, Ontario, Canada, L8N 3Z5.

## Abstract

Hyperinsulinemia can be a cause and consequence of obesity and insulin resistance. Increased insulin secretion and reduced insulin clearance can contribute to hyperinsulinemia. The triggers for changes in insulin clearance during obesity are ill-defined. We found that oral antibiotics mitigated impaired insulin clearance in mice fed a high fat diet (HFD) for 12 weeks or longer. Short-term HFD feeding and aging did not alter insulin clearance in mice. Germ-free mice colonized with microbes from HFD-fed mice had impaired insulin clearance, but not C-peptide clearance, and only after mice were colonized for 6 weeks and then HFD-fed. Five bacterial taxa predicted >90% of the variance in insulin clearance. Our data indicate that gut microbes are an independent and transmissible factor that regulates obesity-induced changes in insulin clearance. A small cluster of microbes may be a target for mitigating defects in insulin clearance and the progression of obesity and Type 2 Diabetes. We propose that a small community in the gut microbiota can impair insulin clearance and increase insulin load and the risk of complications from hyperinsulinemia.

## Introduction

Obesity is a predictor for insulin resistance and increased blood glucose and risk factor for Type 2 Diabetes (T2D). Hyperinsulinemia has been implicated in the progression of obesity, insulin resistance and T2D. Elevated insulin can be a cause and consequence of obesity and insulin resistance[1–3]. It is not yet clear how environmental factors, including gut-resident microbes, alter the relationship between hyperinsulinemia and obesity or insulin resistance. Dynamic insulin responses are controlled by insulin secretion versus insulin clearance coupled with insulin degradation. Increased insulin secretion and reduced (i.e. impaired) insulin clearance can contribute to hyperinsulinemia. Obesity is associated with higher insulin secretion that can occur irrespective of changes in insulin sensitivity, whereas impaired insulin clearance is associated with insulin resistance during obesity[4].

Insulin secretion is widely investigated in obesity and Type 2 diabetes. Pancreatic beta cells sense blood glucose and secrete insulin, which promotes glucose uptake and lipogenesis and inhibits lipolysis and gluconeogenesis. Pancreatic beta cell characteristics and insulin secretion are modulated by neuronal and hormonal inputs and defects in beta cell function underpin the risk of Type 2 Diabetes[5–7]. Insulin clearance is less studied, but clearance of insulin is also be modified by hormones such as incretins. For example, lower insulin clearance can lead to increased blood insulin levels due to glucagon-like peptide-1 (GLP-1) administration in mice[8]. Insulin clearance dynamics can be divided into hepatic and peripheral contributions. After insulin is secreted into the portal vein, insulin initially encounters the liver before accessing the general circulation. Approximately 50-80% of insulin may be depleted from the blood by hepatic uptake and degradation during first-pass insulin clearance[9,10]. Subsequently, skeletal muscle and the kidneys are key tissues that clear blood insulin via tissue-mediated insulin uptake and enzymatic degradation, which can protect against excessive insulin load and hypoglycemia[10].

Pancreatic-derived proinsulin is cleaved into two peptides: the active insulin hormone and C-peptide. Measuring both blood insulin and C-peptide together can estimate the contributions of insulin secretion versus insulin clearance[11]. C-peptide is not subject to the same stringent clearance mechanism of blood insulin, and it is possible to take advantage of this divergence in the mechanisms of hormone clearance to determine the specificity of insulin clearance versus the disappearance of co-secreted C-peptide or general mechanisms of clearance for other peptides.

Insulin clearance is a key regulator of circulating insulin levels[12]. Hepatic insulin clearance is involved in the integrated response regulating insulin sensitivity, glucose production, and lipogenesis[12]. Impaired insulin clearance has been proposed as a contributor to (rather than a consequence of) insulin resistance[13]. Reduced insulin clearance may be driven by impaired hepatic or peripheral clearance, but it is not yet clear how obesity versus insulin resistance influences hepatic or peripheral insulin clearance[4,14]. In obese patients assessed for insulin resistance, the magnitude of lower insulin clearance coincided with a progressive increase in levels of blood insulin[4]. Furthermore, reduced insulin clearance can occur prior to compensatory increases in insulin secretion, suggesting that reduced insulin clearance may be an early physiological response that is integrated into changes in insulin sensitivity[4]. Aging is associated with increased insulin levels and insulin resistance, but a comparison between mice aged 3 and 10 months suggested that hyperinsulinemia associated with this period of aging is related to increased insulin secretion and not reduced insulin clearance[15].

The triggers of impaired insulin clearance during obesity are ill-defined. Obesity is associated with metabolic endotoxemia and lipopolysaccharides (LPS) derived from the cell wall of Gram-negative bacteria can impair insulin clearance[16,17]. Microbial pathogens such as *Salmonella typhimurium* lower insulin clearance and promote insulin resistance in mice[16]. It is already known that the intestinal microbiota can regulate glucose metabolism and insulin secretion. For example, gut microbes can modulate insulin secretion in germ-free mice colonized with the intestinal microbiota of various mouse strains[18]. We hypothesized that gut microbes also regulate insulin clearance, which could contribute to postprandial hyperinsulinemia during diet-induced obesity. Here, we define a role for the intestinal microbiota in regulating insulin clearance during prolonged diet-induced obesity in mice. We found that diet-induced changes in a small number of related taxa can explain the majority of microbe-induced changes in insulin clearance. We found that microbes from obese mice are an independent and transmissible contributor to impaired insulin clearance during diet-induced obesity, which may contribute to hyperinsulinemia, insulin resistance, and obesity.

## Results

### High fat feeding impairs insulin clearance during an oral glucose challenge in mice

Intestinal microbiota can regulate blood glucose and insulin secretion[18,19], but it was unknown if gut microbes regulate insulin clearance. We fed mice an obesogenic, low fiber, HFD or control (chow) diet for 14 weeks. Some mice were supplemented with antibiotics (1 g/L ampicillin and 0.5 g/L neomycin) in the drinking water during the last 2 weeks of high fat feeding. Mice fed HFD had higher body mass and higher fasting blood glucose relative to chow diet-fed mice (Figure 1A, B). Treatment of HFD-fed mice with antibiotics lowered fasting blood glucose (Figure 1B) but antibiotics did not alter body mass (Figure 1A). After 14 weeks on each diet, we performed an oral glucose challenge (4 g/kg, p.o.) and collected blood samples for analysis of insulin and C-peptide plasma concentrations. HFD-fed mice had higher fasting insulin and a greater increase in blood insulin concentration during the oral glucose challenge (Figure 1C). C-peptide was elevated both in the fasted state and during the oral glucose challenge in HFD-fed mice compared to chow diet-fed mice (Figure 1D). Antibiotic treatment attenuated the increase in blood insulin, but not C-peptide, during the oral glucose challenge (Figure 1C, D). Antibiotic treatment also lowered fasting insulin, but not fasting C-peptide levels in the serum (Figure 1C, D). These results suggest that antibiotics improve (i.e. increase) insulin clearance, but do not alter C-peptide kinetics after an oral glucose challenge in mice chronically fed an obesogenic HFD.

**Figure 1:**
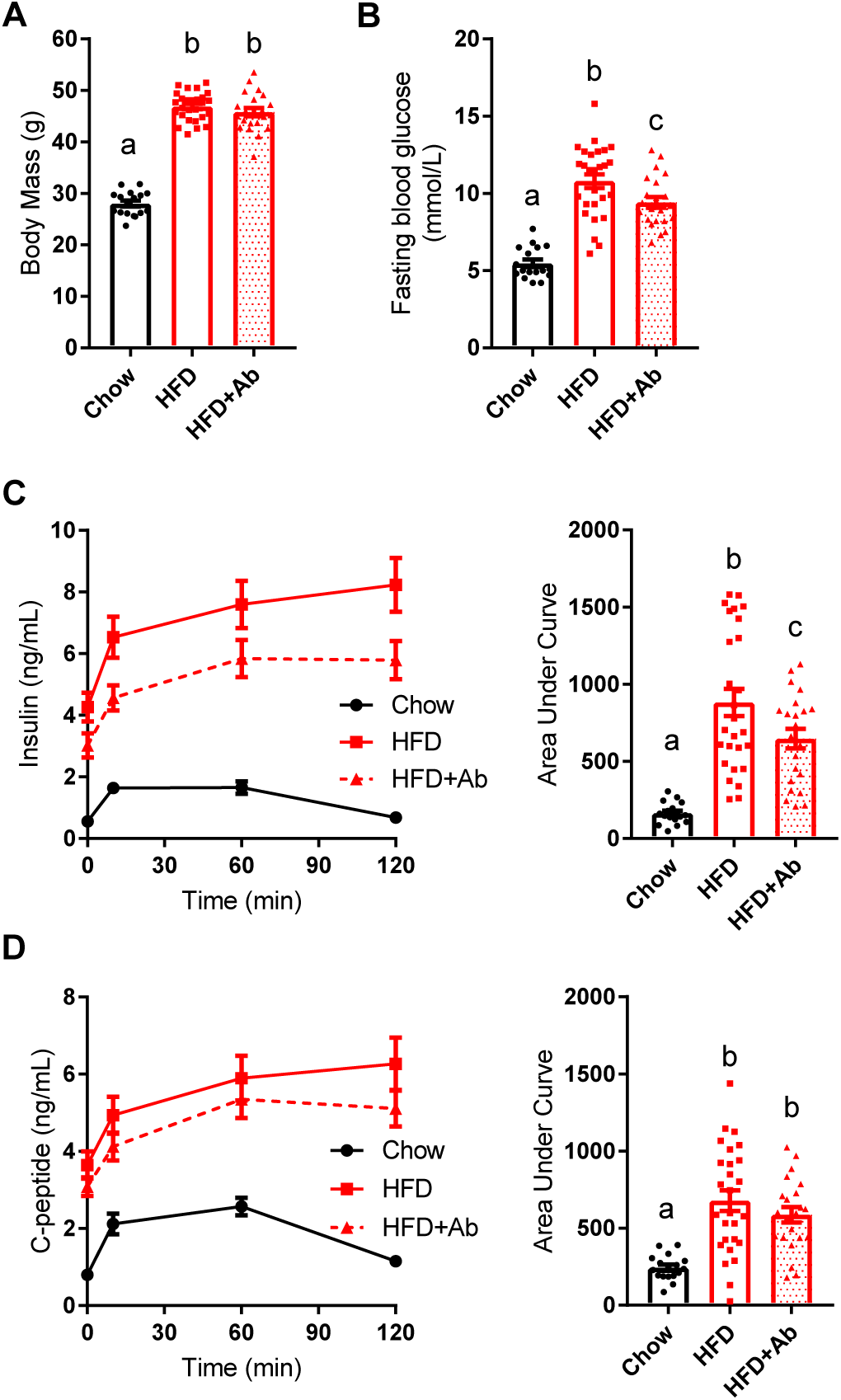
Antibiotics mitigate impaired insulin clearance during an oral glucose challenge in obese mice. Male mice were fed a control (chow) diet or an obesogenic low fibre HFD for 14 weeks. A subset of HFD-fed mice was given antibiotics (1 g/L ampicillin and 0.5 g/L neomycin) in their drinking water during the last 2 weeks (Chow=17, HFD=27, HFD+Ab=22). **A)** Body mass and **B)** fasting blood glucose after week 14 week. **C)** Insulin and **D)** C-peptide were measured in plasma by ELISA, collected from each time point (0, 10, 60, 120 minutes) during an oral glucose challenge (4 g/kg). **Data information:** All values are mean +/-SEM. Statistical significance was measured as p < 0.05 using one-way ANOVA. Post hoc analysis was performed using Tukey’s multiple comparisons test. Groups of mice denoted by different letters are statistically different from one another. Each dot/symbol indicates one mouse.

### Antibiotics rescue impaired insulin clearance caused by long-term high fat feeding

Next, we chose to investigate insulin clearance directly, independently of control mechanisms engaged by an oral glucose load. To this end, human insulin was injected (1 U/kg, i.p.) in lean and obese mice and its presence in circulation was assessed as a readout of insulin clearance. Two weeks of HFD-feeding increased body mass (Figure 2A) but did not alter insulin clearance (Figure 2B). Oral antibiotics did not alter body mass or insulin clearance in lean mice or mice fed HFD for 2 weeks (Figure 2A, B). However, in mice fed HFD for 12 weeks, body mass was increased (Figure 2C) and insulin clearance was lower given the increased area under the curve (AUC) in the plasma insulin concentration versus time (Figure 2D), relative to age-matched chow diet-fed mice. Oral antibiotics during the last 2 weeks of HFD-feeding increased insulin clearance without changing body mass (Figure 2C, D). Similarly, 2 weeks of oral antibiotics increased insulin clearance in mice fed HFD for 37 weeks without changes in body mass (Figure 2E, F). Together, these data support a model in which the intestinal microbiota changes during chronic diet-induced obesity contributes to impaired insulin clearance.

**Figure 2:**
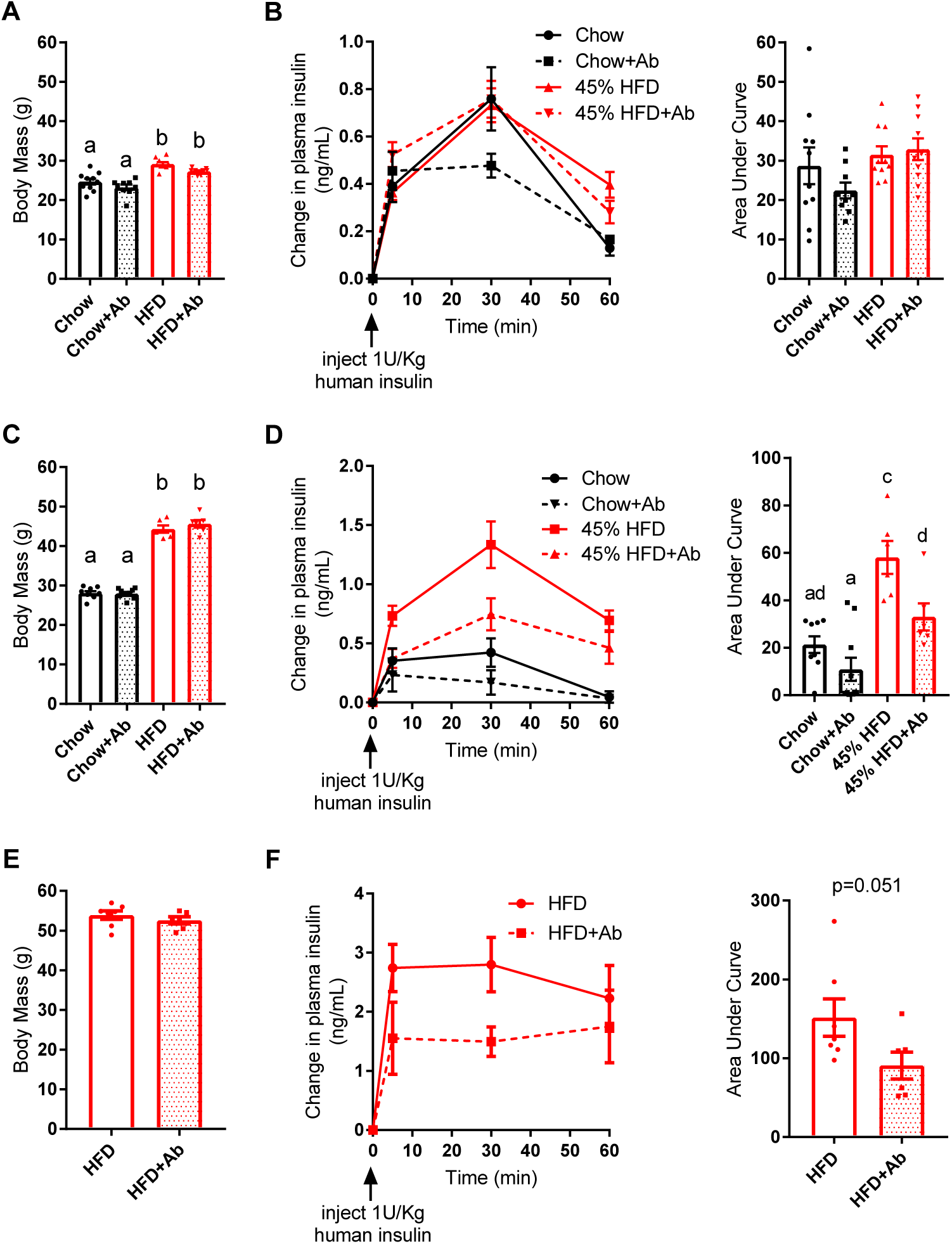
Antibiotics mitigate impaired insulin clearance after chronic HFD feeding in obese mice. **A)** Body mass and **B)** insulin clearance were measured in male mice fed a control chow diet or obesogenic low fiber HFD for 2 weeks +/-antibiotics (1 g/L ampicillin and 0.5 g/L neomycin) in their drinking water (N=9-10 per group). Concentration of human insulin was measured in plasma at indicated time points following injection of human insulin (1U/kg, i.p.). **C)** Body mass and **D)** insulin clearance in male mice fed a chow diet or HFD for 12 weeks followed by 2 additional weeks +/-antibiotics in their drinking water (N=6-10 per group). **E)** Body mass and **F)** insulin clearance in male mice fed HFD for 35 weeks followed by 2 additional weeks +/-antibiotics in their drinking water (N=6-7 per group). **Data information:** All values are mean +/-SEM. Statistical significance was measured as p < 0.05 using two-way ANOVA and Post hoc analysis using Tukey’s multiple comparisons test (A-D) or Student unpaired t-test (E-F). Groups denoted by different letters are statistically different from one another. Each dot/symbol indicates one mouse.

### Insulin Clearance is not altered by aging

Our data show that antibiotics improve insulin clearance in mice fed HFD for a prolonged period, such as 12 or 37 weeks, but not during short-term HFD feeding (i.e. 2 weeks), irrespective of increased body mass. Aging alters the composition of the gut microbiome and promotes inflammation and insulin resistance[20]. To investigate if aging contributes to defects in insulin clearance, we tested insulin clearance in chow-fed mice that were 4, 10, or 21 months of age. Although body mass increased with age (Figure 3A), insulin clearance was unaltered in aged mice (Figure 3B). Thus, our data support a model where defects in insulin clearance in mice fed a HFD for 12 weeks or longer occurred independently of aging. These data are concordant with a previous report that demonstrated aging did not alter insulin clearance in 3- and 10-month-old mice[15].

**Figure 3:**
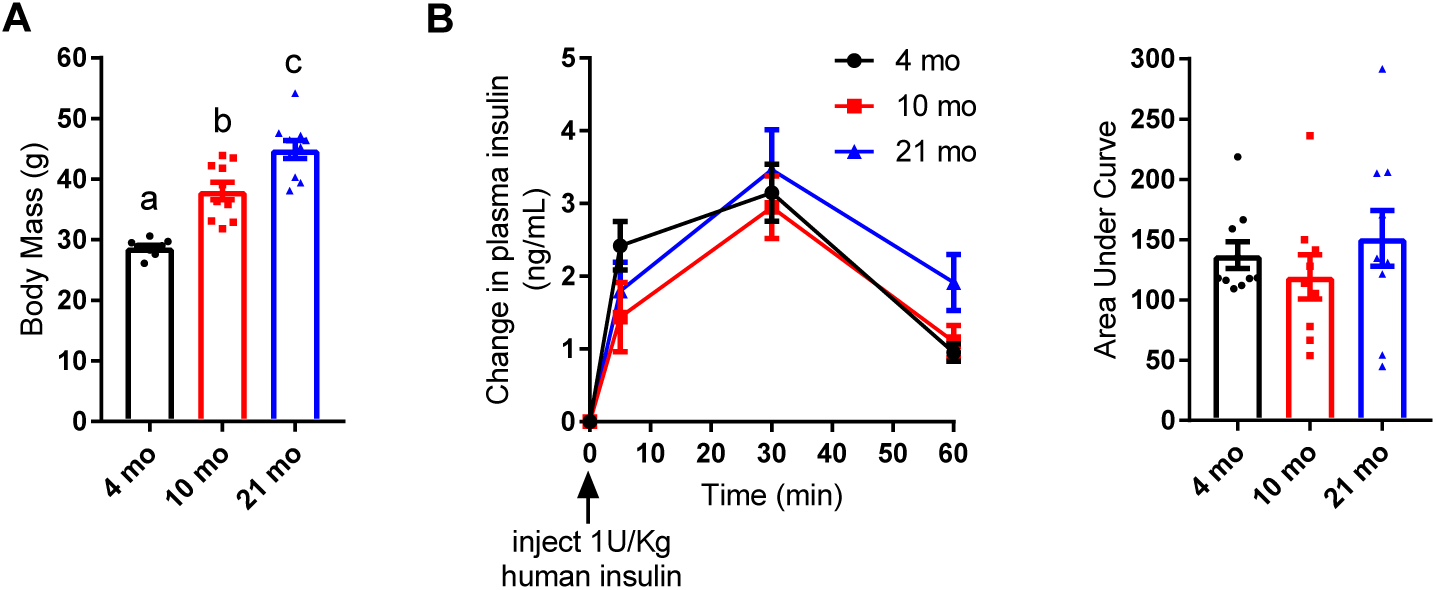
Insulin clearance is not altered across the life course of mice. Male mice were fed a control chow diet for 4 months, 10 months, or 21 months (N=10 per group). **A)** Body mass **B)** Insulin clearance after injection of human insulin (1U/kg, i.p.). **Data information:** All values are mean +/-SEM. Statistical significance was measured as p < 0.05 using one-way ANOVA. Post hoc analysis was performed using Tukey’s multiple comparisons test. Groups denoted by different letters are statistically different from one another. Each dot/symbol indicates one mouse.

### The microbiota from HFD-fed mice is an independent and transmissible factor that impairs insulin clearance

To investigate a causative role for the intestinal microbiota in regulating insulin clearance, we colonized germ-free mice for 6-8 weeks with the microbiota from conventional donor mice fed either a chow or HFD containing 45% of calories from fat (Figure 4A). Following 6-8 weeks of continual colonization, all recipient mice were switched from the chow diet to the HFD. The germ-free mice colonized with microbes from HFD-fed donors (HFD-R mice) had higher body mass than those mice colonized with microbes from chow-fed donors (Chow-R mice), when assessed after 6 weeks on chow diet and after 2 weeks of HFD feeding (Figure 4B).

**Figure 4:**
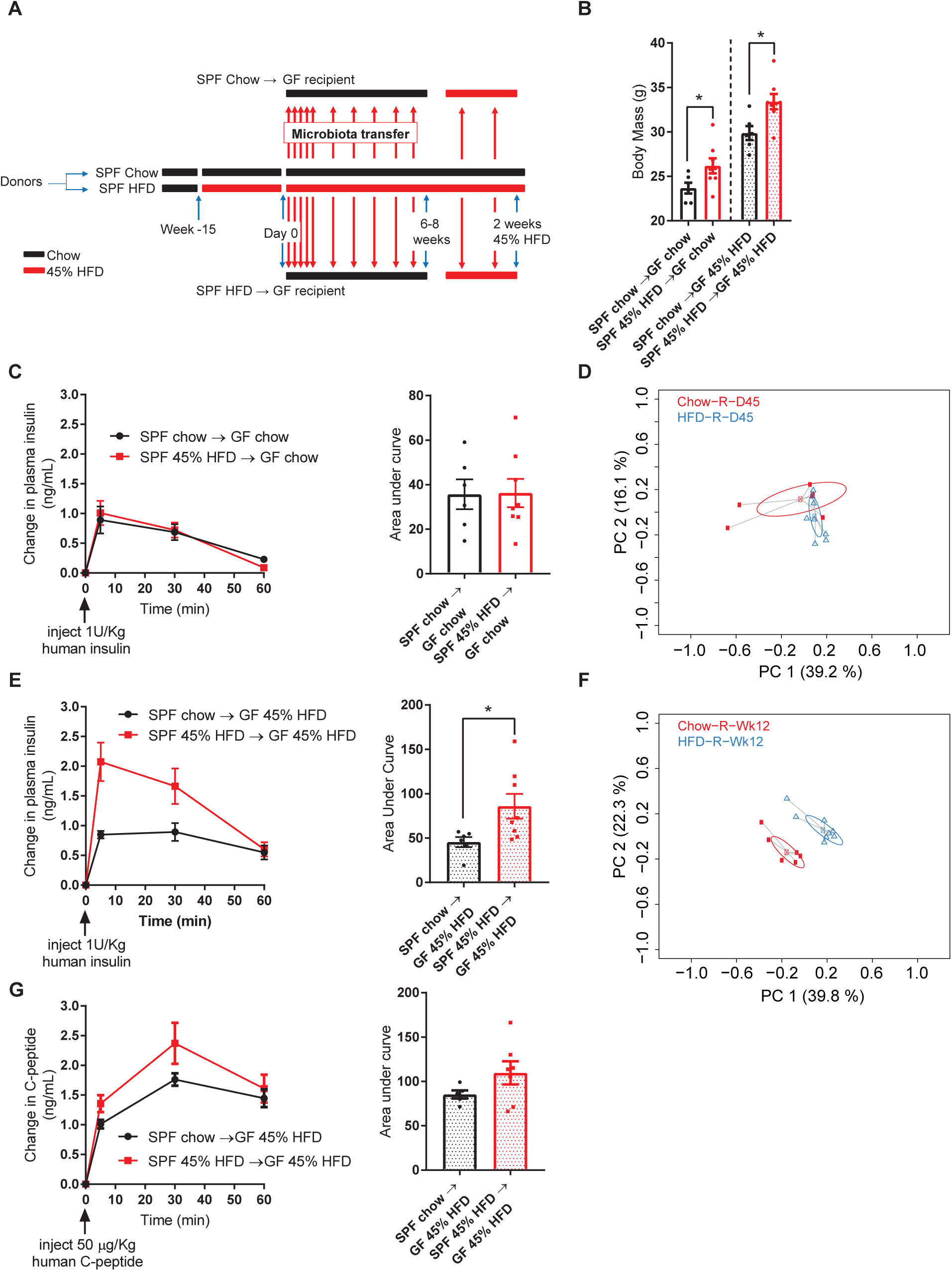
Gut microbiota from HFD-fed mice is an independent and transmissible factor for impaired insulin clearance. **A)** Schematic of experimental design where specific pathogen free (SPF) donor mice were placed on control chow diet or and obesogenic low fiber (45%) HFD for 15 weeks. On Day 0, and each subsequent day, germ-free mice (“recipients”) were colonized with the microbiota from the donor mice for 7 days and then microbial colonization was reinforced once per week for 8-10 weeks. All germ-free recipient mice were on chow diet for 6-8 weeks then all germ-free recipient mice were switched from a chow diet to HFD for 2 additional weeks. **B)** Body mass of germ-free recipient mice colonized with microbiota from chow diet fed mice or HFD-fed mice where recipient mice were assessed after 6 weeks on a chow diet or after 2 weeks of HFD feeding (N=6-8). **C)** Insulin clearance after injection of human insulin (1U/kg, i.p.) was measured in germ-free recipient mice after 6 weeks of colonization, where all recipient mice were fed chow diet (N=6-8). **D)** PCoA plot of Bray-Curtis dissimilarity of the gut microbiota from germ-free recipient mice at the time of the insulin clearance test in (C). **E)** Insulin clearance after injection of human insulin (1U/kg, i.p.) measured in germ-free recipient mice after 8 weeks of colonization on a chow diet plus an additional 2 weeks on HFD. **F)** PCoA plot of Bray-Curtis dissimilarity of the microbiota from germ-free recipient mice at the time of the insulin clearance test in (E). **G)** C-peptide clearance after injection of human C-peptide (50 µg/kg) clearance in germ-free recipient mice after 6 weeks of colonization on a chow diet plus an additional 2 weeks on HFD (N=5-7). **Data information:** All values are mean +/-SEM. Statistical significance was measured as p < 0.05 using Student t-test. Each dot/symbol indicates one mouse.

On day 45 of colonization, germ-free recipient mice were challenged with human insulin (1 U/kg) and tested for insulin clearance (Figure 4C). At this time, all recipient mice were being fed chow diet. There was no difference in insulin clearance between Chow-R and HFD-R mice (Figure 4C). Surprisingly, there was also no significant difference in the microbial composition between Chow-R and HFD-R mice fed a chow diet, as measured by the Bray-Curtis dissimilarity index (Figure 4D). We have previously used this 45-day colonization protocol to cause transmissible differences in the intestinal microbiota and glucose metabolism of germ-free recipient mice colonized with microbes, but donor mice were fed a more obesogenic HFD that contain 60% of calories from fat[19]. In the current experiments used donor mice fed a HFD with 45% of calories derived from fat and the absence of an overt microbial phenotype may explain why there was no difference in insulin clearance. It was also possible that subtle differences in the microbiota existed between the Chow-R and HFD-R groups, but a dietary stress in the recent mice was required to unmask the phenotype. To test this hypothesis, we switched all recipient mice from a chow diet to HFD for 2 weeks and repeated the human insulin (1 U/kg) challenge. Based on the data in Figure 2B, which show that 2 weeks of HFD feeding is insufficient to cause defective insulin clearance, we predicted that this intervention should not alter insulin clearance unless an underlying microbial phenotype was present. The Chow-R group displayed an almost identical insulin clearance rate after 2 weeks on HFD compared to chow diet feeding (Figure 4E), consistent with our data in Figure 2B. However, the HFD-R mice now (after 2 weeks of HFD feeding) displayed impaired insulin clearance relative to the Chow-R group (Figure 4E). These data suggest that the microbes in HFD-R predisposed mice towards defective insulin clearance, which manifested upon the stress of HFD feeding. Importantly, this transmissible microbiota effect on insulin clearance was unlikely due to differences in body mass between Chow-R and HFD-R mice, as these groups differed in body mass both on chow diet and HFD (Figure 4B). β-diversity analysis confirmed that the microbial populations of Chow-R and HFD-R mice were different after 2 weeks of HFD feeding (Figure 4F). Thus, the stress of HFD feeding revealed differences in the composition of microbiota of Chow-R versus HFD-R mice, which resulted in defective insulin clearance in the HFD-R mice. We also tested C-peptide clearance in colonized germ-free Chow-R and HFD-R mice after 2 weeks of HFD using a human C-peptide injection (50 µg/kg, i.p.). C-peptide clearance was not altered in HFD-R relative to Chow-R, demonstrating that the microbiota of HFD-fed mice does not alter clearance of a protein co-secreted with insulin (Figure 4G). Our findings show that the microbiota of lean mice is sufficient to protect mice from obesity-induced microbe-transmissible defects in insulin clearance. Our findings also indicate that microbial and dietary factors synergize to impair insulin clearance.

### HFD-induced dysbiosis is associated with defective insulin clearance

To identify microbial taxa that correlate with impaired insulin clearance during diet-induced obesity, we compared the fecal microbiome characteristics in Chow-R and HFD-R mice at day 45 (Figure 4D) and after 2 weeks of HFD feeding (Figure 4F) using a pairwise Wilcoxon test of all taxa (collapsed to the genus level). We hypothesized that select taxa would be the uniquely changed in HFD-R mice after 2 weeks of high fat feeding and that these taxa would be key microbial predictors of impaired insulin clearance. The relative abundances of 32 operational taxonomic units (OTUs) significantly differed between Chow-R and HFD-R, on either day 45 or after 2 weeks of HFD feeding (Figure 5A).

**Figure 5:**
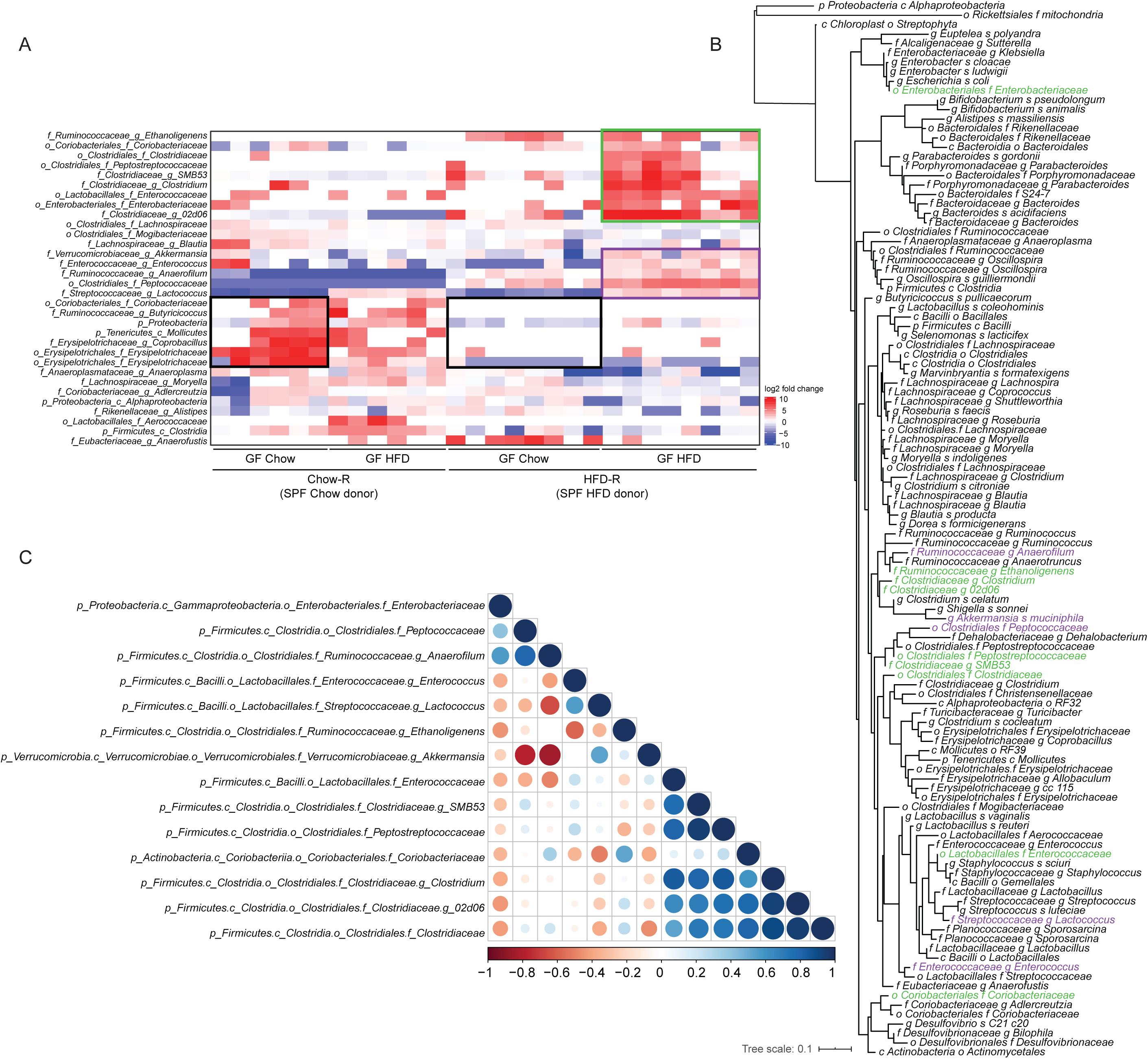
A cluster of phylogenetically related microbes correlate with impaired insulin clearance. **A)** Clustered heatmap of the 32 microbial taxa that were significantly different between germ-free recipient mice colonized with microbiota from mice fed a control chow diet versus mice fed an obesogenic, low fiber HFD (total 28 samples). Recipient mice colonized with microbes were compared after 6 weeks on chow diet and then after 2 weeks of HFD feeding. Fold change in the relative abundance of each taxon was calculated relative to the median level across the 28 samples and plotted in the heatmap. Three clusters of taxa were identified based on differences between Chow-R and HFD-R groups (cluster 1 = 2 black boxes, cluster 2 = purple box, cluster 3 = green box on the heatmap). Statistical analysis was performed using pairwise Wilcoxon rank sum test. Correction for multiple hypothesis testing (FDR) was calculated using the Benjamini-Hochberg method. Statistical significance was accepted at *p* < 0.05. **B)** Phylogenetic tree of the 16S rRNA genomic sequences of all the taxa detected at a minimum of 10 reads in all the recipient germ-free recipient mice. Bacterial taxa from clusters 2 and 3 are highlighted in purple and green, respectively. **C)** Pearson co-correlation analysis of the taxa from clusters 2 and 3 of the heatmap in panel A in order to identify pairs that are closely correlated in their relative abundance.

Three clusters of taxa can be delineated in the feces of Chow-R and HFD-R mice. First, a cluster of 7 taxa shows lower relative abundance in HFD-R mice versus Chow-R mice, when both groups of mice were fed chow diet (see Figure 5A – 2 black boxes on the heat map). These lower abundances persisted when both recipient groups were switched to HFD. Since HFD-R mice did not display defective insulin clearance when fed chow diet (Figure 4C), these differences are unlikely to account for the changes in insulin clearance. The second cluster shows a group of 5 taxa whose relative abundances were only increased when HFD-R mice were fed HFD (see Figure 5A – purple box on the heat map). These taxa show small to moderate differences between Chow-R and HFD-R fed chow diet, but these taxa were clearly increased in HFD-R mice fed HFD. The third cluster shows the most dramatic shift in relative abundances related to defective insulin clearance (see Figure 5A – green box on the heat map). These 9 taxa were very low or below detectable levels in the Chow-R group, irrespective of diet. In the HFD-R group, these taxa were low or absent when mice were fed chow diet, but these 9 taxa significantly increased in relative abundance when HFD-R mice were switched to HFD, which is the condition that coincides with impaired insulin clearance.

To understand the phylogenetic relationship between the taxa identified in these clusters, we created a phylogenetic tree of the 16S rRNA genomic sequences for all taxa detected in the recipient germ-free mice (Figure 5B, and enlarged circular dendrogram in Supplemental Figure 1). Most taxa from clusters 2 and 3, highlighted in purple and green, respectively, are closely phylogenetically related. Four members of cluster 3 are from the *Clostridiaceae* family and 6 of its 9 members cluster tightly. One member of cluster 3, from the *Enterococcaceae* family, shares similarity with two members of cluster 2. Of the 14 taxa highlighted in clusters 2 and 3, only two members, from the *Enterobacteriaceae* and *Coriobacteriaceae* family, are located on more distal nodes. Notably, no bacterial taxa significantly decreased with the emergence of the insulin clearance defect. Changes in taxa associated with cluster 1 are not good candidates in the search for microbes that could drive defective insulin clearance, as the levels of these taxa were comparable across the HFD-fed Chow-R and HFD-R mice. However, microbes identified in clusters 2 and 3, which are phylogenetically similar, are potential candidates for driving defective insulin clearance during diet-induced obesity.

### A consortium of five bacterial taxa predicts changes in insulin clearance

We next performed Pearson correlations of the relative abundance of each taxon from clusters 2 and 3 (see Figure 5A) to the AUC from the insulin clearance test performed in Figure 4E (Supplemental Figure 2). We focused only on the germ-free mice colonized with HFD microbiota (HFD-R) after 2 weeks of HFD feeding because this is the condition that had impaired insulin clearance relative to Chow-R mice. In this way, a positive correlation indicates that increased relative abundance occurs in concert with decreased (i.e. worse) insulin clearance. Nine out of 14 taxa positively correlated with degree of insulin clearance (Table 1), although only two of the independent correlations were statistically significant. Table 1 lists the taxa in order of appearance in Figure 5A and ranks taxa 1-14 in order of highest to lowest correlation coefficients. The top 7 correlations were contained in the third cluster, with 4 of these taxa belonging to the family *Clostridiaceae*. These correlations suggest that the taxa contained in cluster 3 are the best candidates for driving microbe-induced defects in insulin clearance.

**Table 1:** Pearson correlations for microbial taxa versus AUC insulin clearance.

| Taxa | R | p-value | Rank |
| --- | --- | --- | --- |
| p_Firmicutes.c_Clostridia.o_Clostridiales.f_Ruminococcaceae.g_Ethanoligenens | 0.203 | 0.63 | 9 |
| p_Actinobacteria.c_Coriobacteriia.o_Coriobacteriales.f_Coriobacteriaceae | 0.38 | 0.352 | 7 |
| p_Firmicutes.c_Clostridia.o_Clostridiales.f_Clostridiaceae | 0.514 | 0.193 | 4 |
| p_Firmicutes.c_Clostridia.o_Clostridiales.f_Peptostreptococcaceae | 0.428 | 0.29 | 5 |
| p_Firmicutes.c_Clostridia.o_Clostridiales.f_Clostridiaceae.g_SMB53 | 0.405 | 0.319 | 6 |
| p_Firmicutes.c_Clostridia.o_Clostridiales.f_Clostridiaceae.g_Clostridium | 0.764 | **0.027** | 2 |
| p_Firmicutes.c_Bacilli.o_Lactobacillales.f_Enterococcaceae | 0.811 | **0.015** | 1 |
| p_Proteobacteria.c_Gammaproteobacteria.o_Enterobacteriales.f_Enterobacteriaceae | -0.334 | 0.418 | 12 |
| p_Firmicutes.c_Clostridia.o_Clostridiales.f_Clostridiaceae.g_02d06 | 0.644 | 0.085 | 3 |
| p_Verrucomicrobia.c_Verrucomicrobiae.o_Verrucomicrobiales.f_Verrucomicrobiaceae.g_Akkermansia | 0.292 | 0.483 | 8 |
| p_Firmicutes.c_Bacilli.o_Lactobacillales.f_Enterococcaceae.g_Enterococcus | -0.168 | 0.692 | 10 |
| p_Firmicutes.c_Clostridia.o_Clostridiales.f_Ruminococcaceae.g_Anaerofilum | -0.427 | 0.291 | 13 |
| p_Firmicutes.c_Clostridia.o_Clostridiales.f_Peptococcaceae | -0.601 | 0.115 | 14 |
| p_Firmicutes.c_Bacilli.o_Lactobacillales.f_Streptococcaceae.g_Lactococcus | -0.246 | 0.558 | 11 |

To identify the minimum consortium of microbes that can predict impaired (i.e. lower) insulin clearance, we performed a multiple linear regression of the taxa relative abundance versus insulin clearance in HFD-R mice after 2 weeks of HFD feeding. Input into the model was based on taxa correlation rank from Table 1. Relative abundances of each taxon were added to the model from the highest to the lowest correlate. Before input into the multiple regression model, we performed a Pearson co-correlation analysis of the relative abundance of the 14 taxa to identify pairs that are colinear (Figure 5C). A strong positive correlation existed between the abundances of 3 members of *Clostridiaceae* (f) taxa that are ranked 2, 3, and 4 in Table 1. Thus, the relative abundances of these 3 taxa were averaged to create a single input into the multiple linear regression model. The model that best explained the variance in insulin clearance in the HFD-R, high HFD-fed, mice contained the top 5 taxa from the Pearson correlations: *Enterococcaceae* (f), 3 members of *Clostridiaceae* (f), and *Peptostreptococcaceae* (f) (Table 2). In this case, 92% of the variance in the insulin clearance AUC was explained by these 5 taxa. Thus, a small consortium of taxa that was selectively increased in the HFD-R mice fed HFD for 2 weeks explains the variance in insulin clearance. These 5 candidate taxa represent the candidate microbial community that constitute an independent and transmissible factor in altering insulin clearance during diet-induced obesity.

**Table 2:** Multiple linear regression model of insulin clearance.

| Dependent variable: AUC insulin clearance |  |  |
| --- | --- | --- |
| Independent variables | Adjusted R<sup>2</sup> | p-value |
| Model 1: p_Firmicutes.c_Bacilli.o_Lactobacillales.f_Enterococcaceae | 0.6004 | 0.0146 |
| Model 2: average of the co-correlated clostridia (f_Clostridiaceae + g_Clostridium + g_02d06) | 0.3188 | 0.08415 |
| Model 3: p_Firmicutes.c_Clostridia.o_Clostridiales.f_Peptostreptococcaceae | 0.04711 | 0.29 |
| Multiple linear regression: Model 1 and Model 2 | 0.556 | 0.05665 |
| Multiple linear regression: Model 1 and Model 2 and Model 3 | 0.9243 | 0.003 |

## Discussion

Our data show that the community of intestinal microbes from mice fed an obesogenic diet is an independent and transmissible factor that regulates insulin clearance in mice. We found that a small cluster of phylogenetically related bacteria, including *Enterococcaceae, Clostridiaceae* and *Peptostreptococcaceae* can explain over 90% of the variance in host insulin clearance after microbial transfer from diet-induced obese mice. Microbe-induced changes in insulin clearance did not alter the kinetics of C-peptide levels, indicating that the microbiota induced an effect specific to insulin rather than a co-secreted endocrine factor or general effect on peptide clearance.

Gut microbiome composition and host genetics have been shown to alter insulin secretion. Consistent with our results, where we found that 3 members of the *Clostridiaceae* family correlated with changes in insulin clearance, previous work that focused on insulin secretion found that *Clostridiaceae* showed the highest correlation with insulin levels[18]. It appears that changes in the relative abundance of members of the *Clostridiaceae* family are positioned to alter insulin dynamics in the host. We also found that increased *Enterococcaceae* were part of the small cluster of taxa that corelated with impaired insulin secretion. This result is consistent changes in *Enterococcaceae* regulating insulin, since intermittent fasting lowers blood insulin and glucose and improved insulin sensitivity, coincident with a decreased relative abundance of *Enterococcaceae* in obese, diabetic, db/db mice[21]. Furthermore, while bacterial LPS has been shown to impair insulin clearance, we have recently shown that members of the *Enterococcaceae* family compartmentalize in the tissues of individuals with T2D, independently of obesity[22]. Altogether, these findings position members of the *Enterobacteriaceae* family as key players in diet-induced dysmetabolism in the host. Given the early onset of defective insulin clearance in the progression to T2D, it appears worthwhile to investigate bacterial strains within *Enterobacteriaeae* that could impair the insulin clearance.

We also found that an increased relative abundance of *Peptostreptococcaceae* was in the cluster of related taxa that corelated with impaired insulin secretion. Others have found that feeding mice a diabetogenic HFD for at least a month increased the relative abundance of *Peptostreptococcaceae*, which was linked to lower upper gut Th17 responses including lower RORγt CD4 T-cells that provide protective immunity from diet-induced insulin resistance[23]. Intriguingly, symbiotic treatment that lowers insulin resistance and blood glucose also lowered the relative abundance of *Peptostreptococcaceae*[23]. Our data add microbe-specific regulation of insulin clearance to the growing body of evidence that specific microbes influence dynamic insulin and glucose responses and these endocrine and metabolic responses are altered by changes in the host-microbe relationship during obesity.

To the best of our knowledge these data are the first to show that microbes from obese mice can transmit defects in insulin clearance, where a small cluster of physiologically related microbes can account for the majority of variance in insulin clearance. There are many reports of associations between gut microbes, obesity and insulin sensitivity or blood glucose[24–26]. For example, we have previously shown that the relative abundance of intestinal *Clostridiaceae* is higher in mice with impaired glucose tolerance due to an obesogenic diet[19]. We also showed that *Clostridiaceae* were higher in mice with microbially-driven glucose intolerance and *Clostridium* was one of 9 significantly different taxa in germ-free mice colonized with microbes from HFD-fed mice[19]. This is consistent with the results here showing elevated *Clostridium* associating with impaired insulin clearance. Others have shown that *Prevotella copri* and *Bacteroides vulgatus* have been identified as key species in the human microbiome that can alter insulin sensitivity and intermediates such as branched-chain amino acids[27]. We have not yet identified the microbial or host metabolites that influence the mechanisms of hepatic or peripheral insulin clearance. An important future goal is to define how changes in the abundance of specific microbial species and their metabolites alter insulin receptor endocytosis of extracellular insulin, Carcinoembryonic antigen-related cell adhesion molecule (CEACAM)-mediated insulin clearance and Insulin-degrading enzyme (IDE) activity in specific tissues. It will also be important to interrogate how specific microbial components alter the mechanism of tissue-specific insulin clearance. It is known that components of the bacterial cell wall, such as LPS and muropeptides, can engage innate immune receptors in the pancreatic environment to alter insulin secretion and peripheral insulin sensitivity[16,28–30]. It is possible that shared microbe-host responses potentiate insulin secretion and impair insulin clearance, which could increase the insulin load over time and increase the risk of complications from hyperinsulinemia, including obesity and insulin resistance.

## Materials and Methods

### Mice

All procedures were approved by McMaster University Animal Ethics Review Board. Specific pathogen free (SPF) *C57BL/6J* mice were born at McMaster University. At 8-12 weeks of age, littermate mice were randomly placed on chow or 45% HFD diets. The control (chow) diet contains 17% calories from fat and ∼13% fiber content (Teklad 22/5 diet, catalogue #8640) and the 45% HFD contains ∼6% fibre content, 45% calories are derived from fat, and the energy density is 4.7 kcal per gram of food (Research Diets, D12451). When indicated, an antibiotic cocktail (1.0 mg/mL ampicillin and 0.5 mg/mL neomycin) was provided in the drinking water and changed every 2 days. Germ-free *C57BL/6N* mice, supplied by the Farncombe Gnotobiotic Unit of McMaster University, were exported at 10-12 weeks of age and immediately colonized using soiled litter from SPF *C57BL/6J* donor mice. Colonization was re-enforced each day for the first week and once per week thereafter using new soiled litter from SPF *C57BL/6J* donor mice[19]. Mice were individually housed using ventilated racks and handled only in the level II biosafety hood[31]. Colonized, previously germ-free mice are referred to as recipient mice and were fed chow diet upon export and maintained on chow diet until switched to HFD, when indicated.

### Insulin clearance

All metabolic tests were performed after 6 h of fasting[32]. For insulin clearance during an oral glucose challenge, fasting blood glucose and blood samples (50 µL) were collected from the tail vein after 6 hours of fasting. Mice were then given a 4 g/kg glucose dose by oral gavage and subsequent blood samples (50 µL) were collected from the tail vein at 10, 60, and 120 minutes post-gavage. For insulin clearance during an insulin challenge, mice were given human insulin (1 U/kg, NovoRapid) or human C-peptide (50 µg/kg, Sigma) by intraperitoneal injection and blood samples were collected by tail vein sampling at 0, 5, 30, and 60 minutes post-injection. All blood samples were kept on ice after collection and then centrifuged at 10,000 g for 10 min at 4°C. Plasma was collected into fresh tubes and stored at −80°C. Mouse insulin and C-peptide were detected by multiplex ELISA (Millipore) kit in the plasma samples collected during the oral glucose challenge. Human insulin (Mercodia) and human C-peptide (Millipore) were detected by ELISA kits in the plasma samples collected during the human insulin or human C-peptide challenges, respectively.

### Bacterial profiling

Fecal samples were collected and processed as described[19]. Briefly, DNA was purified using ZymoBIOMICS DNA kits (Zymo Research Corporation: D4300), but following mechanical disruption, we also conducted 2 enzymatic lysis steps consisting of lysis solution 1 (50 mg/mL lysozyme and 20% RNase – Sigma R6148) at 37°C for 1 hour and lysis solution 2 (25 µL of 25% SDS, 25 µL of 5M NaCl, and 50 µL of 10 mg per mL Proteinase K) at 60°C for 30 min. Illumina compatible PCR amplification of the variable 3 (V3) region of the 16S rRNA gene was completed on each sample before sequencing on the Illumina MiSeq platform. A minimum of 26,000 reads per sample was acquired. Sequenced data was processed using a custom pipeline, Operational Taxonomic Units (OTUs) were grouped using Abundant OTU+ based on 97% similarity, and the 2013 version of the Greengenes reference database was used to assigned taxonomy to OTUs Ribosomal Database Project (RDP) classifier in Quantitative Insights Into Microbial Ecology (QIIME)[19]. QIIME and R scripts were used to generate plots of taxonomy data and to perform statistical tests. Microbial taxonomy was expressed as relative abundance per sample. In heatmaps, relative abundance was expressed as log_2_ fold change from the median of the entire cohort, as described in each figure. All relative abundance values of 0 were assigned 1×10^−7^ in heat maps, the lowest detectable decimal value in the relative abundance, in order to allow the logarithmic transformation of the fold change. Statistical analyses were performed on relative abundance values. R packages used for data analysis and visualization included vegan, ggplot2, tidyr, dplyr, ggtree, and corrplot.

Phylogenetic analysis of the 16S rRNA genomic sequences was performed with QIIME 2 (Bolyen et al. 2019). For every consensus lineage (taxonomic classification) assigned using the QIIME 2 classifier, those that were present at 10 reads or more across all the recipient mice (28 samples) were used for the analysis. For each one of these consensus lineages, the amplicon sequence variant (ASV) that had the highest total number of reads in the dataset was used as a representative of the taxon. A total of 112 sequences were aligned and used to construct a phylogeny using the QIIME 2 - align-to-tree-mafft-fasttree command (Katoh et al. 2002, Price et al. 2010). The phylogenetic tree was edited using the R package ggtree and visualized using the Interactive Tree Of Life (iTOL)[33].

## Statistical analysis

For measurement of insulin or C-peptide during host metabolic tests an unpaired, two-tailed Student’s t-test was used to compare two groups and ANOVA and Tukey’s post hoc analysis was used to compare more than two groups. Statistical significance was accepted at p<0.05. Analysis and data visualization of microbial populations was conducted in R[34]. A Pairwise Wilcoxon test was used for the non-parametric analysis of variance between groups with the significance threshold set to p<0.05. Adjustment for the false discovery rate (FDR) was calculated with the Benjamini-Hochberg method and statistical significance was accepted at p<0.05[35].

## Acknowledgements

This work was supported by a Foundation grant (FDN-154295) from the Canadian Institutes of Health Research (CIHR) to JDS. CYC holds a Farncombe family graduate student scholarship. JFC holds a Farncombe Family postdoctoral fellowship. TCL holds a CIHR doctoral scholarship. FFA holds a CIHR postdoctoral fellowship and Diabetes Canada incentive funding. JDS holds a Canada Research Chair in Metabolic Inflammation.

## Author contributions

KPF researched the data, contributed to design and discussion, and wrote the manuscript. SZ provided all metagenomics analysis and contributed to discussion. BMD, NGB, FFA, JFC, BDH, CYC, MH, and TCL researched the data. JDS researched the data, derived the hypothesis, wrote the manuscript, and is the guarantor of this work.

## The paper explained

### Problem

Obesity is a leading cause of Type 2 Diabetes. Progression from prediabetes to Type 2 Diabetes during obesity is characterized by hyperinsulinemia and insulin resistance. Hyperinsulinemia can be a cause and consequence of insulin resistance. Dynamic insulin responses are controlled by insulin secretion and insulin clearance coupled with insulin degradation. Increased insulin secretion and reduced (i.e. impaired) insulin clearance can contribute to elevated blood insulin. An increased insulin load over time can exacerbate obesity and insulin resistance. While the control of insulin secretion is widely studied in the context of obesity and diabetes, the triggers for changes in insulin clearance during obesity are ill-defined. The intestinal microbiota can regulate glucose metabolism and insulin secretion, but the contribution of gut microbes to the regulation of insulin clearance was unknown.

### Results

We demonstrate that intestinal microbes regulate insulin clearance during diet-induced obesity in mice. Mice fed high fat diet (HFD) for >12 weeks showed impaired insulin clearance, which was partly rescued with oral antibiotics for 2 weeks. This defect in insulin clearance was not observed after only 2 weeks of HFD-feeding, suggesting that microbes affect insulin clearance during protracted obesity. Germ-free mice colonized with microbes from HFD-fed mice had impaired insulin clearance, but not C-peptide clearance, and only after mice were colonized for 6 weeks and then HFD-fed. A small cluster of phylogenetically related bacteria could explain over 90% of the variance in host insulin clearance after microbial transfer from diet-induced obese mice.

### Impact

To the best of our knowledge these data are the first to show that microbes from obese mice can transmit defects in insulin clearance, where a small cluster of physiologically related microbes can account for the majority of variance in insulin clearance. It is possible that shared microbe-host responses potentiate insulin secretion and impair insulin clearance, which could increase the insulin load over time and increase the risk of complications from hyperinsulinemia, including obesity and insulin resistance.

## Data and Code Availability

The datasets generated during the current study are available from the corresponding author on reasonable request. Figures that have associated raw data are 4 and 5. The custom R scripts used for data analysis are available from the corresponding author on reasonable request.

## Figure Legends

**Supplemental Figure 1: Phylogenetic relationship between microbes detected in germ-free recipient mice**.

An enlarged circular dendrogram of the data in Figure 5B.

**Supplemental Figure 2: Correlations between taxa abundance and insulin clearance**. Pearson correlations were generated between the relative of abundance values of each taxon identified in clusters 2 and 3 (x-axis) to the insulin AUC during the insulin clearance test in the HFD-R mice colonized for 6 weeks on a control chow diet and then fed an obesogenic, low fiber HFD diet for 2 weeks (y-axis). The rank order for highest to lowest correlation coefficients are indicated in the top right corner of each plot.

## References

1. Page MM, Johnson JD (2018) Mild Suppression of Hyperinsulinemia to Treat Obesity and Insulin Resistance. Trends Endocrinol Metab 29: 389–399.

2. Shanik MH, Xu Y, Škrha J, Dankner R, Zick Y, Roth J (2008) Insulin Resistance and Hyperinsulinemia. Diabetes Care 31: S262 LP–S268.

3. Mehran AE, Templeman NM, Brigidi GS, Lim GE, Chu K-Y, Hu X, Botezelli JD, Asadi A, Hoffman BG, Kieffer TJ, et al. (2012) Hyperinsulinemia Drives Diet-Induced Obesity Independently of Brain Insulin Production. Cell Metab 16: 723–737.

4. Jung S-H, Jung C-H, Reaven GM, Kim SH (2018) Adapting to insulin resistance in obesity: role of insulin secretion and clearance. Diabetologia 61: 681–687.

5. MacDonald PE, El-kholy W, Riedel MJ, Salapatek AMF, Light PE, Wheeler MB (2002) The Multiple Actions of GLP-1 on the Process of Glucose-Stimulated Insulin Secretion. Diabetes 51: S434 LP–S442.

6. Yamamoto J, Imai J, Izumi T, Takahashi H, Kawana Y, Takahashi K, Kodama S, Kaneko K, Gao J, Uno K, et al. (2017) Neuronal signals regulate obesity induced β-cell proliferation by FoxM1 dependent mechanism. Nat Commun 8: 1930.

7. Song Y, Yeung E, Liu A, VanderWeele TJ, Chen L, Lu C, Liu C, Schisterman EF, Ning Y, Zhang C (2012) Pancreatic beta-cell function and type 2 diabetes risk: quantify the causal effect using a Mendelian randomization approach based on meta-analyses. Hum Mol Genet 21: 5010–5018.

8. Ahrén B, Thomaseth K, Pacini G (2005) Reduced insulin clearance contributes to the increased insulin levels after administration of glucagon-like peptide 1 in mice. Diabetologia 48: 2140–2146.

9. Tokarz VL, MacDonald PE, Klip A (2018) The cell biology of systemic insulin function. J Cell Biol 217: 2273–2289.

10. Najjar SM, Perdomo G (2019) Hepatic Insulin Clearance: Mechanism and Physiology. Physiology 34: 198–215.

11. Tura A, Ludvik B, Nolan JJ, Pacini G, Thomaseth K (2001) Insulin and C-peptide secretion and kinetics in humans: direct and model-based measurements during OGTT. Am J Physiol Metab 281: E966–E974.

12. Bojsen-Møller KN, Lundsgaard A-M, Madsbad S, Kiens B, Holst JJ (2018) Hepatic Insulin Clearance in Regulation of Systemic Insulin Concentrations—Role of Carbohydrate and Energy Availability. Diabetes 67: 2129 LP–2136.

13. Watada H, Tamura Y (2017) Impaired insulin clearance as a cause rather than a consequence of insulin resistance. J Diabetes Investig 8: 723–725.

14. Ohashi K, Fujii M, Uda S, Kubota H, Komada H, Sakaguchi K, Ogawa W, Kuroda S (2018) Increase in hepatic and decrease in peripheral insulin clearance characterize abnormal temporal patterns of serum insulin in diabetic subjects. npj Syst Biol Appl 4: 14.

15. Kurauti MA, Ferreira SM, Soares GM, Vettorazzi JF, Carneiro EM, Boschero AC, Costa-Júnior JM (2019) Hyperinsulinemia is associated with increasing insulin secretion but not with decreasing insulin clearance in an age-related metabolic dysfunction mice model. J Cell Physiol 234: 9802–9809.

16. Hagar JA, Edin ML, Lih FB, Thurlow LR, Koller BH, Cairns BA, Zeldin DC, Miao EA (2017) Lipopolysaccharide Potentiates Insulin-Driven Hypoglycemic Shock. J Immunol 199: 3634–3643.

17. Amar J, Burcelin R, Ruidavets JB, Cani PD, Fauvel J, Alessi MC, Chamontin B, Ferriéres J (2008) Energy intake is associated with endotoxemia in apparently healthy men. Am J Clin Nutr 87: 1219–1223.

18. Kreznar JH, Keller MP, Traeger LL, Rabaglia ME, Schueler KL, Stapleton DS, Zhao W, Vivas EI, Yandell BS, Broman AT, et al. (2017) Host Genotype and Gut Microbiome Modulate Insulin Secretion and Diet-Induced Metabolic Phenotypes. Cell Rep 18: 1739–1750.

19. Foley KP, Zlitni S, Denou E, Duggan BM, Chan RW, Stearns JC, Schertzer JD (2018) Long term but not short term exposure to obesity related microbiota promotes host insulin resistance. Nat Commun 9: 4681.

20. Thevaranjan N, Puchta A, Schulz C, Naidoo A, Szamosi JC, Verschoor CP, Loukov D, Schenck LP, Jury J, Foley KP, et al. (2017) Age-Associated Microbial Dysbiosis Promotes Intestinal Permeability, Systemic Inflammation, and Macrophage Dysfunction. Cell Host Microbe 21: 455–466.e4.

21. Liu Z, Dai X, Zhang H, Shi R, Hui Y, Jin X, Zhang W, Wang L, Wang Q, Wang D, et al. (2020) Gut microbiota mediates intermittent-fasting alleviation of diabetes-induced cognitive impairment. Nat Commun 11: 855.

22. Anhê FF, Jensen BAH, Varin T V, Servant F, Van Blerk S, Richard D, Marceau S, Surette M, Biertho L, Lelouvier B, et al. (2020) Type 2 diabetes influences bacterial tissue compartmentalisation in human obesity. Nat Metab 2: 233–242.

23. Garidou L, Pomié C, Klopp P, Waget A, Charpentier J, Aloulou M, Giry A, Serino M, Stenman L, Lahtinen S, et al. (2015) The Gut Microbiota Regulates Intestinal CD4 T Cells Expressing RORγt and Controls Metabolic Disease. Cell Metab 22: 100–112.

24. Cavallari JF, Schertzer JD (2017) Intestinal Microbiota Contributes to Energy Balance, Metabolic Inflammation, and Insulin Resistance in Obesity. J Obes Metab Syndr 26: 161–171.

25. Anhê FF, Barra NG, Schertzer JD (2020) Glucose alters the symbiotic relationships between gut microbiota and host physiology. Am J Physiol Endocrinol Metab 318: E111–E116.

26. Everard A, Cani PD (2013) Diabetes, obesity and gut microbiota. Best Pract Res Clin Gastroenterol 27: 73–83.

27. Pedersen HK, Gudmundsdottir V, Nielsen HB, Hyotylainen T, Nielsen T, Jensen BAH, Forslund K, Hildebrand F, Prifti E, Falony G, et al. (2016) Human gut microbes impact host serum metabolome and insulin sensitivity. Nature 535: 376.

28. Zhang Q, Pan Y, Zeng B, Zheng X, Wang H, Shen X, Li H, Jiang Q, Zhao J, Meng Z-X, et al. (2019) Intestinal lysozyme liberates Nod1 ligands from microbes to direct insulin trafficking in pancreatic beta cells. Cell Res 29: 516–532.

29. Schertzer JD, Tamrakar AK, Magalhães JG, Pereira S, Bilan PJ, Fullerton MD, Liu Z, Steinberg GR, Giacca A, Philpott DJ, et al. (2011) NOD1 Activators Link Innate Immunity to Insulin Resistance. Diabetes 60: 2206–2215.

30. Cani PD, Amar J, Iglesias MA, Poggi M, Knauf C, Bastelica D, Neyrinck AM, Fava F, Tuohy KM, Chabo C, et al. (2007) Metabolic endotoxemia initiated obesity and insulin resistance. Diabetes 56: 1761–1772.

31. Denou E, Lolmède K, Garidou L, Pomie C, Chabo C, Lau TC, Fullerton MD, Nigro G, Zakaroff-Girard A, Luche E, et al. (2015) Defective NOD 2 peptidoglycan sensing promotes diet-induced inflammation, dysbiosis, and insulin resistance. EMBO Mol Med 7: 259–274.

32. Schertzer JD, Antonescu CN, Bilan PJ, Jain S, Huang X, Liu Z, Bonen A, Klip A (2009) A Transgenic Mouse Model to Study Glucose Transporter 4myc Regulation in Skeletal Muscle. Endocrinology 150: 1935–1940.

33. Letunic I, Bork P (2019) Interactive Tree Of Life (iTOL) v4: recent updates and new developments. Nucleic Acids Res 47: W256–W259.

34. Lex A, Gehlenborg N, Strobelt H, Vuillemot R, Pfister H (2014) UpSet: Visualization of Intersecting Sets. IEEE Trans Vis Comput Graph 20: 1983–1992.

35. Benjamini Y, Hochberg Y (1995) Controlling the False Discovery Rate: A Practical and Powerful Approach to Multiple Testing. J R Stat Soc Ser B 57: 289–300.

